# Maintaining task performance levels under cognitive load while walking requires widespread reallocation of neural resources: A Mobile Brain-Body Imaging (MoBI) study

**DOI:** 10.1101/2023.06.20.545763

**Authors:** Eleni Patelaki, John J. Foxe, Amber L. McFerren, Edward G. Freedman

## Abstract

The neural underpinnings of increasing cognitive load during walking, despite being ubiquitous in everyday life, is still not fully understood. This study elucidates the neural mechanisms underlying increased cognitive load while walking, by employing 2 versions of a Go/NoGo response inhibition task, namely the 1-back Go/NoGo task and the more cognitively demanding 2-back Go/NoGo task, during sitting or walking on a treadmill. By using the Mobile Brain/Body Imaging (MoBI) modality, electroencephalographic (EEG) activity, three-dimensional (3D) gait kinematics and task-related behavioral responses were collected from 34 young adults for the 1-back Go/NoGo task and 34 young adults for the 2-back Go/NoGo task. Interestingly, increasing cognitive-inhibitory load from 1-back to 2-back Go/NoGo during walking was not associated with any detectable costs in response accuracy, response speed, or gait consistency; however, it came with attenuations in walking-related EEG amplitude changes during both successful inhibitions (correct rejections) and successful executions (hits) of the ‘Go’ motor response. During correct rejections, such attenuations were detected over frontal regions, during latencies related to sensory gain control, conflict monitoring and working memory storage and processing. During hits, attenuations were found over left-parietal regions, during latencies related to orienting attention to and selecting the ‘Go’ motor plan, as well as over central regions, during latencies linked to executing the ‘Go’ motor response. The pattern of attenuation in walking-related EEG amplitude changes, manifested by the 2-back Go/NoGo group, is thought to reflect more effortful recalibration of the above neural processes, a mechanism which might be a key driver of performance maintenance in the face of increased cognitive demands while walking. Overall, the present findings shed light on the extent of the neurocognitive capacity of young adults, thus revealing the employed methodology as promising for better understanding how factors such as aging or neurological disorders could impinge on this capacity.

## INTRODUCTION

Successfully engaging in everyday activities requires performance of tasks of varied cognitive demands, from simple tasks such as lifting a cup of tea, to complex ones such as calculating one’s monthly budget. Oftentimes, the environmental demands necessitate the concurrent performance of two or more tasks, known as multitasking. One of the most common multitasking scenarios in daily life is performing a cognitive task while walking (hereafter referred to as dual-task walking), and this has been shown to have various effects on the performance of young adults, ranging from deterioration [1–5], to no detectable impact [6–8], or even improvement [8–10] in terms of response speed and accuracy, or gait speed and variability. Whether dual-task walking has negative, neutral, or positive effects on behavior presumably depends on how the two concurrent tasks interact with each other at a neural-resource level, which, in turn, relates to the difficulty of the walking and cognitive tasks. Most studies pairing disturbance-free walking with relatively simple cognitive tasks in young adults show minimal to no deterioration or even frank improvement in gait and cognitive task performance [6, 7, 11–15]. However, increasing the difficulty of either the walking or the cognitive task has been shown to elicit clearer decrements in young adults, more consistent with the ‘cognitive-motor interference’ hypothesis (CMI) [16, 17]. Specifically, increasing motor load beyond simple walking (e.g. walking in an obstacle course) while engaging in a cognitive task, has been reported to reduce response accuracy [18, 19] and increase gait variability [14]. Similarly, increasing cognitive load through more complex cognitive tasks while walking has been linked to reduced response speed [20], gait speed [1, 4] and response accuracy [1], and increased gait variability [4].

In young adults, increasing cognitive load during walking has been found to elicit changes in frontoparietal functional connectivity [14, 20] and prefrontal blood oxygenation [13, 21], even in the absence of associated behavioral decrements in response accuracy, gait speed or gait variability [13, 14, 20, 21]. These findings suggest that young adults can effectively recruit additional neural resources to compensate for the increase in task demands. However, the nature of these adaptations in neural processing remains unclear. Although oxygenation and connectivity measures can quantify load-related changes in neural resource usage and interplay, they lack the temporal resolution required to examine which specific stages of information processing are affected by increasing cognitive task load during walking. The event-related potential (ERP) technique, on the other hand, can capture neurophysiological changes at a millisecond timescale [22–24] and, as such, it constitutes an appropriate tool for studying the temporal aspect of load-related processing changes while walking.

Response inhibition, which is the ability to suppress prepotent, inappropriate responses to stimuli, thoughts, or emotions, is critical for successfully carrying out everyday tasks [25], and is considered one of the canonical components of executive control processing. When the stimulus, thought, or emotion to which a response needs to be inhibited must be simultaneously rehearsed in working memory, then inhibitory difficulty increases markedly, presumably because response inhibition and working memory compete for the allocation of common neural substrates [26–29]. One well-established approach to studying response inhibition is the visual 1-back Go/NoGo task, which requires pressing a response button after each unique image is presented (‘Go’ trial), but withholding the button press when an image is a repeat of the previous image (1-back ‘NoGo’ trial) [30, 31]. In order to increase the inhibitory load through manipulating the working memory component, a novel integration of tasks is proposed here, which is called the 2-back Go/NoGo task and it differs from the 1-back Go/NoGo task in that the button press now has to be withheld when an image is a repeat of the image preceding the previous image (2-back ‘NoGo’ trial). In the 1-back Go/NoGo task, successfully inhibiting a button press during a ‘NoGo’ trial typically elicits two stimulus-locked ERP components, the N2 and the P3. The N2 is a negative voltage deflection with a frontocentral scalp topography, peaking around 200-350 ms post-stimulus-onset [32–34]. The N2 has been linked to conflict monitoring processes subserved by the anterior cingulate cortex (ACC) [32, 35–37]. The P3 is a positive voltage deflection elicited around 350-600 ms post-stimulus-onset [6, 38], and its scalp topography spans from parietal to frontal regions. During these latencies, both motor and non-motor stages of inhibitory control are implemented [39, 40], which explains why generation of the P3 has been associated with multiple cortical sources [36, 41–46]. Failing to inhibit a button press during a ‘NoGo’ trial typically elicits the error-related negativity (ERN), which is a response-locked ERP component peaking ~50 ms after the erroneous button press [47, 48]. The ERN has been linked to conflict monitoring during errors, and it has a frontocentral scalp distribution which putatively reflects its generation from the ACC and associated frontal regions [35, 49]. The 2-back Go/NoGo task integrates response inhibition with working memory in a novel manner; since it is an inhibitory task, N2s/P3s and ERNs are expected to be elicited here as well during successful and unsuccessful inhibitions, respectively. However, it is still unknown how the additional working memory load will alter these ERP components.

This study aimed to investigate how increasing inhibitory load while walking is managed at a cognitive-systems level. We hypothesized that increasing inhibitory load from 1-back to 2-back Go/NoGo during walking would be marked by a decline in behavioral performance and reallocation of neural resources. To test this hypothesis, load-related changes in inhibitory and gait-kinematic behavior, as well as EEG neural activity, were probed. Studying how the neurocognitive systems of young healthy adults respond to increasing load while walking can help determine their ‘tipping point’, namely the point beyond which the imposed load starts outweighing the available compensatory mechanisms, thus resulting in the manifestation of behavioral performance declines. Identifying the tipping point of young adults and comparing it with the tipping point of older adults, or that of individuals with neurological disorders, can provide a deeper understanding of the trajectories of cognitive decline in these populations.

## MATERIALS AND METHODS

### Participants

Sixty-one (61) young adults participated in the study: 27 performed the 1-back Go/NoGo task only (18-30 years old; age = 22.09 ± 3.12 years; 17 female, 17 male; 30 right-handed, 4 left-handed), 27 performed the 2-back Go/NoGo task only (19-34 years old; age = 22.53 ± 3.69 years; 16 female, 18 male; 30 right-handed, 4 left-handed), and 7 performed both tasks (at different dates). This resulted in 68 datasets in total: 34 datasets for the 1-back Go/NoGo task and 34 datasets for the 2-back Go/NoGo task. For details about the tasks, see the Experimental Design section below.

All participants provided written informed consent, reported no diagnosed neurological disorders, no recent head injuries, and normal or corrected-to-normal vision. The Institutional Review Board of the University of Rochester approved the experimental procedures (STUDY00001952). All procedures were compliant with the principles laid out in the Declaration of Helsinki for the responsible conduct of research. Participants were paid the lab-standard hourly rate for time spent in the lab.

### Experimental Design

Two versions of a Go/NoGo response inhibition cognitive task were employed: a 1-back version and a 2-back version. In both versions, during each experimental block, images were presented in the central visual field for 67 ms with a fixed stimulus-onset-asynchrony of 1017 ms. On average, images subtended 20° horizontally by 16° vertically. The task was coded using the Presentation software (version 20.1, Neurobehavioral Systems, Albany, CA, USA). Participants were instructed to press the button of a wireless game controller using their dominant hand as fast and accurately as possible for every image, except when the presented image was the same as the previous image for the 1-back Go/NoGo task (1-back ‘NoGo’ trial, Fig. 1A), or as the image preceding the previous image for the 2-back Go/NoGo task (2-back ‘NoGo’ trial, Fig. 1B). Participants performed blocks of 240 trials in which 209 (87%) were Go trials and 31 (13%) were NoGo trials. NoGo trials were randomly distributed within each block.

**Fig. 1.**
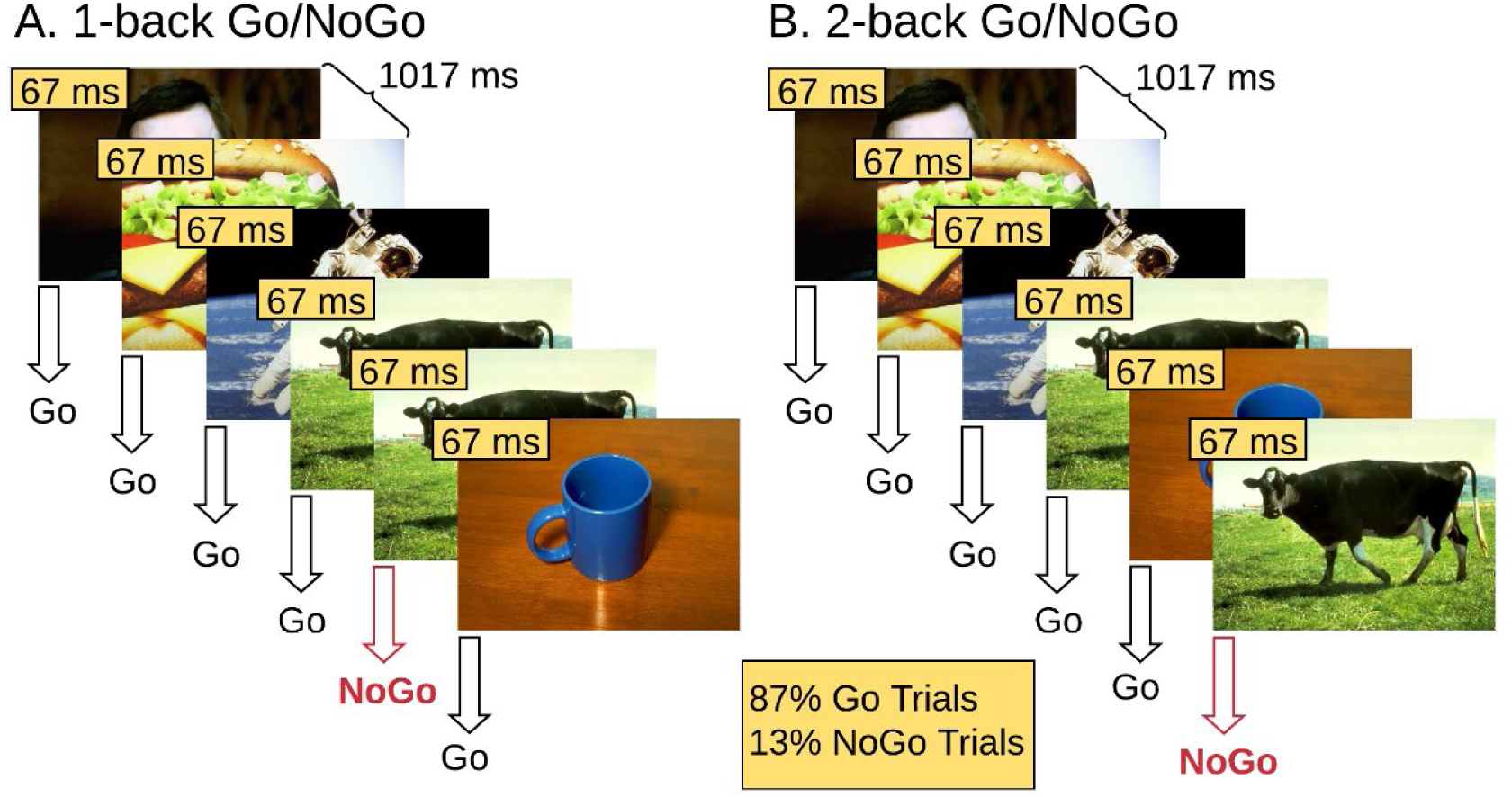
Illustration of the A. 1-back and B. 2-back Go/NoGo response inhibition experimental design. Participants are instructed to respond on Go trials and withhold response on NoGo trials.

The images used for stimuli were drawn from the International Affective Picture System (IAPS) database [50]. The IAPS database contains pictures of varied emotional valence and semantic content. Positive, neutral and negative pictures were all used, however analyzing the emotional valence or semantic content of stimuli is beyond the scope of this study.

Four behavioral conditions of the cognitive task were defined: 1) hits, defined as the Go trials on which participants correctly pressed the response button, 2) correct rejections, defined as the NoGo trials on which participants correctly withheld their response, 3) false alarms, defined as the NoGo trials on which participants incorrectly pressed the response button and 4) misses, defined as the Go trials on which participants failed to press the response button.

For both task versions, experimental blocks were performed while the participants were either sitting or walking on a treadmill (Tuff Tread, Conroe, TX, USA), at a distance of 2.25 m approximately from the projection screen on which the images were projected (Barco F35 AS3D, 1920×1080 pxl). A safety harness was worn while walking to guard against falls (https://youtu.be/HS-5Qk5tvDE). An experimental session consisted of 16 blocks: 1 training block at the beginning, 7 sitting blocks, 7 walking blocks, and a walking-only block (walking on the treadmill without a cognitive task). The order of sitting and walking blocks was pseudorandomized; no more than 3 consecutive walking blocks occurred to prevent fatigue. Participants were allowed to take short breaks between the blocks, each of which lasted 4 minutes. Most participants took at least one break during the experiment. If a break was requested, typically it did not last longer than 10 minutes. Participants were asked to select a treadmill speed corresponding to brisk walking for them. On average, participants walked at 4.64 ± 0.42 km/h for the 1-back Go/NoGo task, and at 4.84 ± 0.23 km/h for the 2-back Go/NoGo task.

EEG data were recorded using a BioSemi Active Two System (BioSemi Inc., Amsterdam, the Netherlands) and a 64-electrode configuration following the International 10-20 system [51]. Neural activity was digitized at 2048 Hz. Full-body motion capture was recorded using a 16 camera OptiTrack system (Prime 41 cameras), and Motive software (OptiTrack, NaturalPoint, Inc., Corvallis, OR, USA) in a ~37 m^2^ space. Cameras recorded 41 markers on standard anatomical landmarks along the torso, the head and both arms, hands, legs and feet at 360 frames per second. Stimulus triggers from Presentation (Neurobehavioral Systems Inc., Berkeley, CA, USA), behavioral responses from the game controller button, motion tracking data and EEG data were time-synchronized using Lab Streaming Layer (LSL) software (Swartz Center for Computational Neuroscience, University of California, San Diego, CA, USA; available at: https://github.com/sccn/labstreaminglayer). Motion capture data were recorded using custom software written to rebroadcast the data from the Motive software to the LSL lab recorder. EEG data were recorded from available LSL streaming plugins for the BioSemi system. Behavioral event markers were recorded using the built-in LSL functionality in the Presentation software. The long-term test-retest reliability of the MoBI approach has been previously detailed [52]. All behavioral, EEG and motion kinematic data processing and basic analyses were performed using custom MATLAB scripts (MathWorks Inc., Natick, MA, USA) and/or functions from EEGLAB [53]. Custom analysis code will be made available on GitHub (https://github.com/CNL-R) upon publication.

### Cognitive Task Performance Processing & Analysis

The timing of each button press relative to stimulus onset, the participant’s response times (RTs), were recorded using the Response Manager functionality of Presentation and stored with precision of 1/10 millisecond. The Response Manager was set to accept responses only after 183 ms post-stimulus-onset within each experimental trial. Any responses prior to that were considered delayed responses to the previous trial and were ignored. This RT threshold was selected to filter out as many delayed-response trials as possible, without rejecting any valid trials for which the responses were merely fast [54].

The behavioral conditions of the cognitive task that were examined in this study in terms of EEG activity were 1) hits, 2) correct rejections, and 3) false alarms. For correct rejections and false alarms, only trials that were preceded by at least 1 hit in the 1-back Go/NoGo task, and at least 2 hits in the 2-back Go/NoGo task, were included. This was done to ensure that the inhibitory component was present. Correct rejections and false alarms were selected based on previous studies reporting walking-related EEG changes during these behavioral conditions [10, 55]. To examine whether the presence or absence of the inhibitory component potentially affected walking-related EEG changes, hits were examined as well.

Two behavioral measures were calculated: 1) the d’ score (sensitivity index) and 2) mean RT during (correct) Go trials, namely hits. D’ is a standardized score and it is computed as the difference between the Gaussian standard scores for the false alarm rate (percentage of unsuccessful NoGo trials) and the hit rate (percentage of successful Go trials) [56, 57]. D’ score was preferred over correct rejection rate (percentage of successful NoGo trials) as a measure of accuracy of inhibitory performance, since it removes the bias introduced by different response strategies adopted across participants. For a more detailed explanation, the reader is referred to the ‘Cognitive task performance processing & analysis’ section of Methods, in Patelaki and colleagues [10].

### EEG Activity Processing & Analysis

EEG signals were first filtered using a zero-phase Chebyshev Type II filter (*filtfilt* function in MATLAB, passband ripple *Apass* = 1 dB, stopband attenuation *Astop* = 65 dB) [58], and subsequently down-sampled from 2048 Hz to 512 Hz. Next, ‘bad’ electrodes were detected based on kurtosis, probability, and the spectrum of the recorded data, setting the threshold to 5 standard deviations of the mean, as well as covariance, with the threshold set to ±3 standard deviations of the mean [58]. These ‘bad’ electrodes were removed and interpolated based on neighboring electrodes, using spherical interpolation. All the electrodes were re-referenced offline to a common average reference.

It has been shown that 1-2 Hz highpass filtered EEG data yield the optimal Independent Component Analysis (ICA) decomposition results in terms of signal-to-noise ratio [59, 60]. In order to both achieve a high-quality ICA decomposition and retain as much low-frequency (< 1 Hz) neural activity as possible, after running Infomax ICA (*runica* function in EEGLAB, the number of retained principal components matched the rank of the EEG data) on 1-45 Hz bandpass-filtered data and obtaining the decomposition matrices (weight and sphere matrices), these matrices were transferred and applied to 0.01-45 Hz bandpass-filtered data. High-pass filtering was selected to be conservative based on evidence that high-pass filters ≤ 0.1 Hz introduce fewer artifacts into the ERP waveforms [61]. ICs were labeled using the ICLabel algorithm [62]. ICs whose sum of probabilities for the five artifactual IC classes (‘Muscle’, ‘Eye’, ‘Heart’, ‘Line Noise’, ‘Channel Noise’) was higher than 50% were labeled as artifacts and were thus rejected. The remaining ICs were back-projected to the sensor space [60, 63, 64].

Subsequently, the resulting neural activity was split into temporal epochs. For both correct rejection and hit trials, epochs were locked to the stimulus onset, beginning 200 ms before and extending until 800 ms after stimulus onset of the trial. Both correct rejection and hit epochs were baseline-corrected relative to the pre-stimulus-onset interval from −100 to 0 ms. For the false alarm trials, epochs were locked to the response onset, beginning 500 ms before and extending until 500 ms after response onset of the trial. False alarm epochs were baselined-corrected relative to the pre-response interval from −400 to −300 ms. Epochs with a maximum voltage greater than ±150 µV or that exceeded 5 standard deviations of the mean in terms of kurtosis and probability were excluded from further analysis. Epochs that deviated from the mean by ±50 dB in the 0-2 Hz frequency window (eye movement detection) and by +25 or −100 dB in the 20-40 Hz frequency window (muscle activity detection) were rejected as well. On average, 26% (27% for 1-back Go/NoGo and 24% for 2-back Go/NoGo) of the trials were rejected for the sitting condition, and 42% (44% for 1-back Go/NoGo and 40% for 2-back Go/NoGo) for the walking condition. Event-related potentials (ERPs) were measured by averaging epochs for (2 motor-task)x(3 cognitive-task) conditions, namely 6 experimental conditions in total. The motor task conditions were 1) sitting and 2) walking; the behavioral conditions of the cognitive task were 1) hits, 2) correct rejections and 3) false alarms. Table 1 shows the number of trials that were accepted for each motor task condition, behavioral condition of the cognitive task, and inhibitory load group, before the averaging step.

**Table 1.** Number of accepted trials per motor task condition, cognitive task condition and inhibitory load group.

|  | 1-back |  | 2-back |  |
| --- | --- | --- | --- | --- |
|  | sitting | walking | sitting | walking |
| hits | 958 ± 191 | 743 ± 299 | 1000 ± 150 | 808 ± 221 |
| correct rejections | 89 ± 39 | 74 ± 40 | 72 ± 26 | 60 ± 27 |
| false alarms | 59 ± 29 | 40 ± 27 | 77 ± 20 | 61 ± 22 |

### Gait Processing & Analysis

Heel markers on each foot were used to track gait kinematics. The three dimensions (3D) of movement were defined as follows: X is the dimension of lateral movement (right-and-left relative to the motion of the treadmill belt), Y is the dimension of vertical movement (up-and-down relative to the motion of the treadmill belt), and Z is the dimension of fore-aft movement (parallel to the motion of the treadmill belt). The heel marker motion in 3D is described by the three time series of the marker position over time in the X, Y and Z dimension, respectively. Gait cycle was defined as the time interval between two consecutive heel strikes of the same foot. Heel strikes were identified as the local maxima of the Z position waveform over time. To ensure that no ‘phantom’ heel strikes were captured, only peaks with a prominence greater than 0.1 m were kept (*findpeaks* function in MATLAB, *minimum peak prominence* parameter was set to 0.1 m).

Stride-to-stride variability was quantified as the mean Euclidean distance between consecutive 3D gait cycle trajectories, using the Dynamic Time Warping algorithm (DTW) [65, 66]. DTW is an algorithm for measuring the similarity between time series, and its efficacy in measuring 3D gait trajectory similarity is well-established [67–69].

In the case of one-dimensional signals, if X_m=1,2,..,M_ the reference signal and Y_n=1,2,..,N_ the test signal, then DTW finds a sequence {ix, iy} of indices (called warping path), such that X(ix) and Y(iy) have the smallest possible distance. The ix and iy are monotonically increasing indices to the elements of signals X, Y respectively, such that elements of these signals can be indexed repeatedly as many times as necessary to expand appropriate portions of the signals and thus achieve the optimal match. This concept can be generalized to multidimensional signals, like the 3D gait cycle trajectories which are of interest here. The minimal distance between the reference and the test signals (gait trajectories here) is given by formula (1):

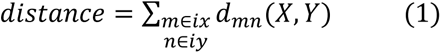

Gait cycle trajectories with a kurtosis that exceeded 5 standard deviations of the mean were rejected as outliers. Also, before DTW computation, gait cycle trajectories were resampled to 100 samples. Since DTW essentially calculates the sum of the Euclidean distances between corresponding points of two interrogated trajectories, ensuring that all trajectories are resampled to the same length helps avoid bias in the algorithm computations.

The actual measure that was used to quantify each participant’s stride-to-stride variability is the mean across DTW distances occurring from all stride-to-stride comparisons. Right-foot and left-foot stride-to-stride DTW distances were pooled to calculate the mean DTW distance per participant.

### Statistical Analyses

Statistical analyses were conducted to test the effects of increasing inhibitory load on cognitive task performance, EEG activity and gait kinematics. All the measures used in the context of these statistical analyses were selected a priori based on previous work from our group [10, 55].

#### Cognitive Task Performance

##### Response Accuracy

To assess how increasing inhibitory load impacts response accuracy during the ‘baseline’ sitting condition as well as walking-related response accuracy change, sitting d’ score and walking-*minus*-sitting d’ score were correlated with inhibitory load. Inhibitory load was defined as a binary variable (2 possible values: 1-back, 2-back). To test the correlation between a binary and a continuous variable, the most appropriate choice is point-biserial correlation, which is mathematically equivalent to Pearson correlation [70]. Since all correlations conducted in the context of this study tested the relationship between a binary variable (namely, inhibitory load) and a continuous physiological or behavioral variable, the Pearson correlation was used for all of them. If walking-*minus*-sitting d’ score was found to significantly correlate with inhibitory load, then d’ score difference between sitting and walking was follow-up tested within both load groups (1-back, 2-back) using paired t-tests in the case of normally distributed data, and Wilcoxon signed rank tests in the case of non-normally distributed data. Follow-up testing aimed to determine whether the potential significant correlation was driven by a specific load group.

To examine the effects of walking on response accuracy at a single-subject level, participants were subsequently classified into three groups, based on whether their d’ score during walking was 1) significantly higher than during sitting (d’walking > d’sitting; response accuracy improved significantly while walking; **IMP**), 2) not statistically different from their d’ score during sitting (d’walking ≈ d’sitting; response accuracy did not change significantly while walking; **NC** = No Change), or 3) significantly lower than during sitting (d’walking < d’sitting; response accuracy declined significantly while walking; **DEC**). The individual walking-*minus*-sitting d’ score of each participant was defined significant if it lay outside of the 95% confidence interval of the normal distribution that had a mean value of zero and a standard deviation equal to that of the walking-*minus*-sitting d’ score distribution of the entire cohort, pooled across load levels. This classification methodology has been proposed and described previously by Patelaki and colleagues [10, 55].

##### Response Speed

Walking-*minus*-sitting mean response time (RT) during hits was correlated with inhibitory load (binary variable, possible values: 1-back, 2-back) using a Pearson correlation, to test for potential relationships between increasing inhibitory load and response speed during walking. Given that previous research has revealed a speed-accuracy trade-off when adding walking to Go/NoGo task performance [55], in order to obtain the true effects of inhibitory loading on response speed during walking, it is important to disentangle them from any effects of response accuracy. To this end, if inhibitory load was found to correlate with any of the d’-related variables defined in the ‘Response Accuracy’ section (walking-*minus*-sitting d’ score, sitting d’ score), then this variable was controlled for by replacing the initial Pearson correlation with a partial Pearson correlation. If the correlation was found to be significant, then mean RT difference between sitting and walking was follow-up tested within both load groups (1-back, 2-back). For follow-up testing, paired t-tests were used if the data were normally distributed, and Wilcoxon signed rank tests if the data were non-normally distributed.

#### EEG Activity

The EEG statistical analyses were performed using the FieldTrip toolbox [71] (http://fieldtriptoolbox.org). As stated in the Introduction, load-related changes in ERP amplitudes during walking were expected to be detected during N2 and P3 latencies during correct rejections (approximately [200, 350] ms and [350, 600] ms, respectively [6, 30, 32–34, 38]), as well as during ERN latencies during response-locked false alarms (approximately [−50, 100] ms [72, 73]. Even though the N2, P3 and ERN are more strongly elicited over midline scalp regions, walking-related changes in voltage amplitudes have been found in lateral regions as well during the latencies of these ERP components [10, 55]. Additionally, walking-related changes have been found during latencies other than the N2, P3 or ERN, such as during the N1 [55]. These observations highlight the need to examine the whole scalp and all possible timepoints for load-related changes in ERP amplitudes during walking, which can be implemented using cluster-based permutation tests [74]. Although this is an inherently exploratory approach, the hypothesis-driven part of this study related to testing the N2, P3 and ERN can also be satisfied under the same statistical analysis [10, 55].

Walking-*minus*-sitting mean ERP amplitudes during hits, correct rejections and false alarms were calculated at each electrode and timepoint for each participant, by subtracting the within-subject mean (across trials) sitting ERP waveform from the within-subject mean walking ERP waveform. Subsequently, for each of the behavioral conditions (hits, correct rejections, false alarms), walking-*minus*-sitting mean ERP amplitude was subjected to Pearson correlation with inhibitory load, at each electrode-timepoint pair, to assess whether increasing inhibitory load impacts ERP amplitudes during walking. Given that previous research has revealed patterns of d’-related ERP amplitude changes during walking [55], in order to obtain the true effects of inhibitory loading on ERP amplitudes during walking, it is crucial to disentangle them from any effects of response accuracy. To this end, if inhibitory load was found to correlate with any of the d’-related variables interrogated above (walking-*minus*-sitting d’ score, sitting d’ score), then this variable was controlled for by replacing the initial Pearson correlation with a partial Pearson correlation, at each electrode-timepoint pair. To identify spatiotemporal clusters of significant correlation effects while accounting for multiplicity of pointwise electrode-timepoint testing, cluster-based permutation tests were performed using the Monte Carlo method (10000 permutations, significance level of the permutation tests *a* = 0.05, probabilities corrected for performing 2-sided tests) and the weighted cluster mass statistic [75] (cluster significance level *a* = 0.05, non-parametric cluster threshold). The results of the pointwise correlations from all 64 electrodes and all timepoints were displayed as a statistical clusterplot, which provides a compact and easily interpretable graphical representation of the magnitude, latency onset/offset, and topography of the detected correlation effects [76, 77]. The x, y, and z axes, respectively, represent time, electrode location, and the t-statistic (indicated by a color value) at each electrode-timepoint pair. The t-statistic value at electrode-timepoint pairs that did not belong to any cluster of significant effects was masked, namely set to zero (corresponding to teal color in the clusterplot).

For Pearson correlations, the t-statistic was calculated based on formula (2) (*r* = correlation coefficient, *N* = sample size) [70, 78]:

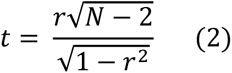

Within each of the interrogated behavioral conditions (hits, correct rejections, false alarms), if any clusters of significant correlation effects were found, then, for each cluster and each of the 2 load groups (1-back, 2-back), mean ERP amplitude, averaged across electrode-timepoints pairs belonging to the cluster, was follow-up tested for differences between sitting and walking. Follow-up tests were performed using paired t-tests if the data were normally distributed, and Wilcoxon signed rank tests if the data were non-normally distributed.

#### Gait

Dual-task-related change in stride-to-stride variability, quantified as WT-*minus*-WO mean DTW distance (WT = walking with task, WO = walking only), was subjected to Pearson correlation with inhibitory load, to test for potential relationships between increasing inhibitory load and stride-to-stride variability when dual tasking. Similar to the EEG and response speed analyses, if inhibitory load was found to correlate with any of the d’-related variables interrogated above (walking-*minus*-sitting d’ score, sitting d’ score), then this variable was controlled for by replacing the initial Pearson correlation with a partial Pearson correlation. If the correlation was found to be significant, then mean DTW distance difference between WO and WT was follow-up tested within both load groups (1-back, 2-back). For follow-up testing, paired t-tests were used if the data were normally distributed, and Wilcoxon signed rank tests if the data were non-normally distributed.

It should be pointed out that one 1-back dataset and one 2-back dataset were excluded from the stride-to-stride (DTW) variability analysis due to lack of walking-only kinematic data, thus resulting in a sample size of 66 datasets (33 1-back, 33 2-back) for this analysis.

## RESULTS

### Cognitive Task Performance

#### Response Accuracy

When engaging in the 1-back Go/NoGo response inhibition task, most young adults have been shown to improve in terms of response accuracy when walking compared to sitting [10]. To assess the impact of increasing inhibitory load under dual-task walking conditions, this study additionally employed the more demanding 2-back Go/NoGo response inhibition task, during both sitting and walking. Regarding the effects of increasing inhibitory load on response accuracy, inhibitory load was found not to correlate with dual-task-related response accuracy change, which was quantified as walking-minus-sitting d’ score (Pearson’s r = −0.04, p = 0.7762). However, inhibitory load was also tested for correlation with sitting d’ score, to investigate any potential effects of increasing inhibitory load on response accuracy independently of walking, and it was found that increased inhibitory load correlated with significantly lower d’ scores during sitting (Pearson’s r = −0.25, p = 0.0421). This finding suggests that, while 2-back Go/NoGo load leads to decline in response accuracy overall, combining it with walking does not impact response accuracy differently than combining walking with the 1-back Go/NoGo task.

Previous work from our group examined at a single-subject level how the d’ performance of young and older adults in the 1-back Go/NoGo task changed between sitting and walking [10, 55]. Here, the same single-subject approach was used to additionally inspect the effects of walking on the d’ performance of young adults when performing the 2-back Go/NoGo task. Fig. 2 shows the distributions of d’ score during sitting and walking, in each of the two inhibitory load groups (1-back, 2-back). As illustrated in Fig. 2, most young adults improved when dual-task walking, both for the 1-back and for the 2-back inhibitory load level. Specifically, from the 34 young adults who performed the 1-back Go/NoGo task, 20 improved when walking (IMPs), 6 did not change significantly across motor load condition (NCs) and 8 declined when walking (DECs). From the 34 young adults who performed the 2-back Go/NoGo task, 20 were classified as IMPs, 8 as NCs and 6 as DECs (Fig. 2). The method used to determine whether d’ change between sitting and walking within each participant was significant or not was the one described in Patelaki and colleagues [10] (for details, see the Statistical Analyses: Cognitive Task Performance: Response accuracy section of the Methods). The finding of improvement in terms of d’ score when young adults are exposed to 1-back Go/NoGo load, though contradictory to the ‘cognitive-motor interference’ (CMI) hypothesis, it is consistent with previous work from our group [10]. Surprisingly, this study revealed that young adults maintain this dual-task-related improvement even when inhibitory load increases from 1-back to 2-back Go/NoGo during walking, which is contrary to our initial hypothesis which predicted response accuracy decline under these dual-tasking conditions.

**Fig. 2.**
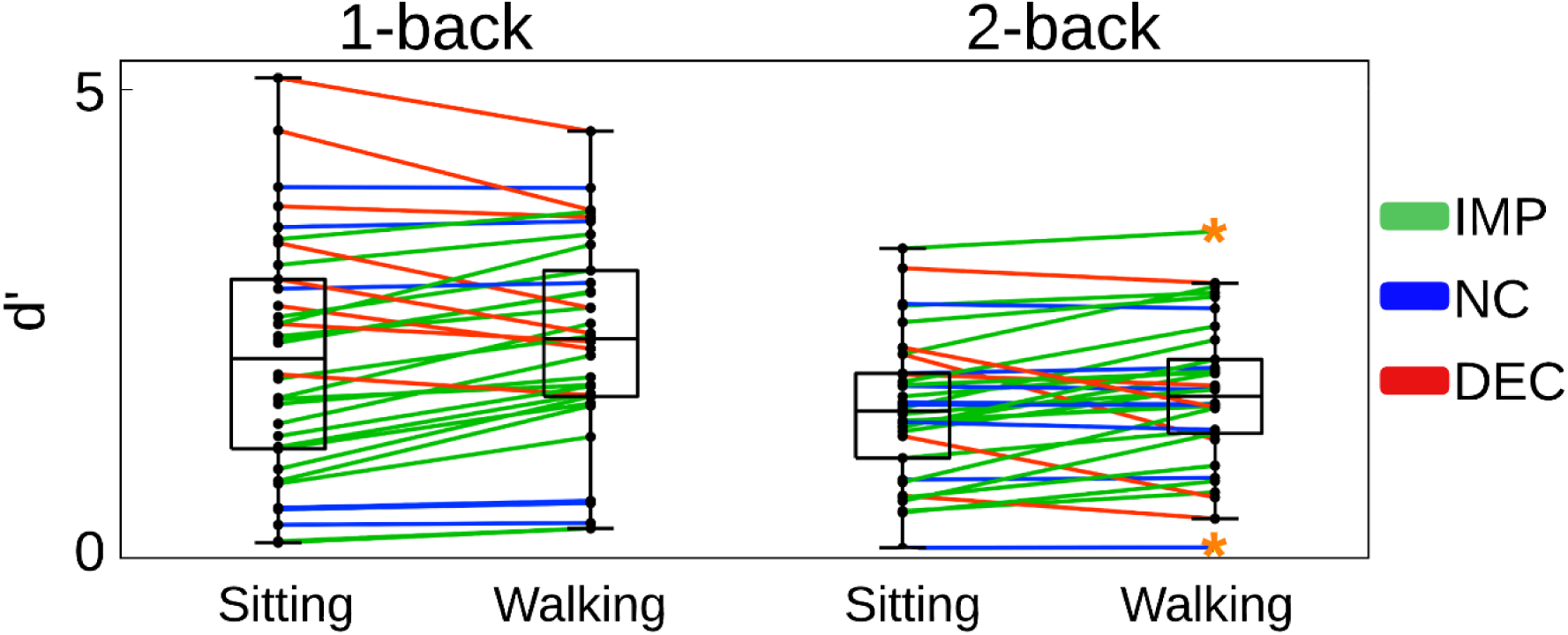
Sitting and walking d’ score distributions, in young adults when engaging in the 1-back and the 2-back Go/NoGo task, respectively. Each line corresponds to one participant. Participants who improved when walking (IMPs; d’walking > d’sitting) are shown in green, those who did not change significantly across motor load condition (NCs; d’walking ≈ d’sitting) are shown in blue, and those who declined when walking (DECs; d’walking > d’sitting) are shown in red. Black dots on the vertical line represent individual participants. The central mark of each box indicates the median, and the bottom and top edges indicate the 25th and 75th percentiles, respectively. The whiskers extend to the most extreme data points not considered outliers. Orange asterisks represent outliers.

Previous studies have shown that d’ performance correlates with ERP activity when walking is combined with a Go/NoGo response inhibition task [10, 55]. Based on these previous findings, in order to obtain the true effects of inhibitory load on dual-task-related ERP amplitude changes, it is important to factor out any d’-related effects. Since increased inhibitory load was found here to correlate with lower d’ scores during sitting, dual-task-related ERP amplitude changes (quantified as walking-*minus*-sitting mean ERP amplitudes) were tested for partial correlation with inhibitory load (1-back, 2-back) while controlling for sitting d’ score. Besides ERPs, the effects of increasing inhibitory load on response speed and stride-to-stride variability during walking were additionally examined, in order to investigate whether this type of increase in dual-task demands was accompanied by costs in other behavioral or physiological domains beyond neural activity. Regarding response speed, since d’ performance has been previously shown to correlate with response speed when walking is added to the employed Go/NoGo task [55], the effects of inhibitory load on dual-task-related response speed change were assessed by subjecting walking-*minus*-sitting mean response time during hits to partial correlation with inhibitory load while controlling for sitting d’ score. Similarly, the effects of inhibitory load on dual-task-related stride-to-stride variability change were assessed by subjecting WT-minus-WO mean DTW distance (WT = walking with task, WO = walking only, DTW = Dynamic Time Warping) to partial correlation with inhibitory load while controlling for sitting d’ score.

#### Response Speed

Fig. 3 shows the distributions of mean response time (RT) during sitting and walking, in each of the two load groups, namely 1-back and 2-back.

**Fig. 3.**
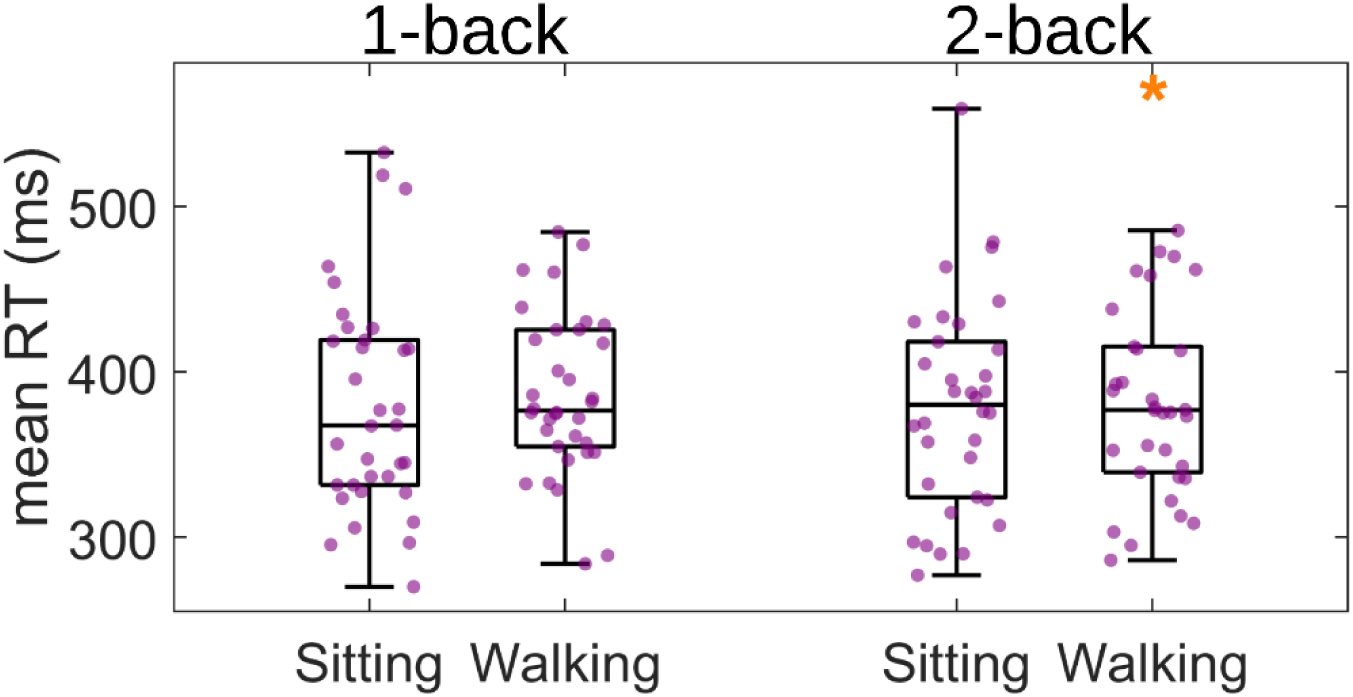
Sitting and walking mean response time (RT) distributions during hits (correct Go trials), in young adults when they engaged in the 1-back and the 2-back Go/NoGo task, respectively. Purple dots represent individual participants. Orange asterisks represent outliers.

Walking-*minus*-sitting mean response time during hits was tested for partial correlation with inhibitory load while controlling for sitting d’ score, using a partial Pearson correlation. This correlation was found to be non-significant (Pearson’s r = −0.08, p = 0.5152), indicating that increase in inhibitory load was accompanied by no detectable cost in response speed when dual tasking.

### EEG Activity

Fig. 4 shows the grand average ERP waveforms during sitting (blue) and walking (red) at three central electrode locations illustrated : FCz, Cz and CPz. Observed ERPs during ‘hits’, correct rejections (‘CRs’) and response-locked false alarms (‘FAs’) are shown for both for 1-back and for 2-back young adults. These electrodes were selected for illustration because the N2 and P3 components during correct rejections and the ERN component during response-locked false alarms have maximal amplitudes over these midline electrode locations [6, 39, 79–81]. For hits and correct rejections, ERP waveforms were aligned on stimulus presentation (vertical line - time = 0) onset, and they were plotted from 200 ms pre-stimulus-onset to 800 ms post-stimulus-onset. The N2 component is the negative voltage deflection spanning approximately from 200 to 350 ms post-stimulus-onset [32–34, 82, 83], and it is evident during both hits and correct rejections. Smaller N2 amplitudes were observed in 2-back (Fig. 4B) compared to 1-back (Fig. 4A) young adults, especially during correct rejections. The P3 component is the positive voltage deflection extending approximately from 350 to 600 ms post-stimulus-onset [6, 38]. In both load groups, P3 was more strongly evoked during correct rejections than during hits, and over centro-parietal topographies (Cz and CPz electrodes), consistent with previous findings in young adults [6, 39]. For false alarms, ERP waveforms were aligned on motor response, namely the button press, (vertical line - time = 0), and they were plotted from −500 ms pre-response to 500 ms post-response. The ERN component is the negative voltage deflection spanning approximately from −50 to 100 ms [72, 73]. ERN was more evident frontocentrally (FCz electrode) in both load groups, and it appeared attenuated in the 2-back compared to the 1-back Go/NoGo load condition. Subsequent analyses will focus on testing walking-related ERP amplitude changes during hits, correct rejections and false alarms, for partial correlation with inhibitory load while controlling for sitting d’ score.

**Fig. 4.**
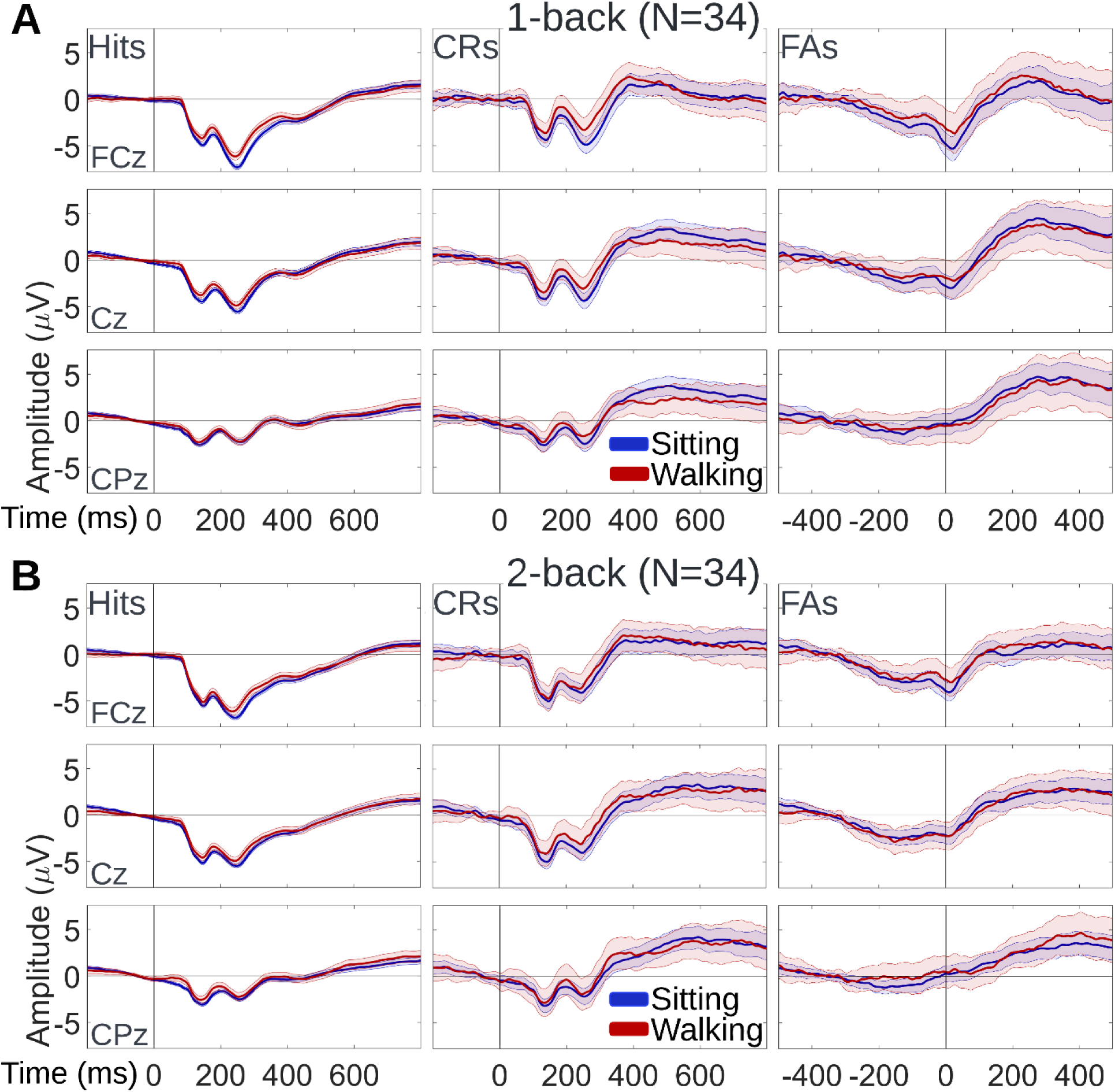
Grand mean sitting (blue) and walking (red) ERP waveforms, at three midline electrode locations: frontocentral midline (FCz), central midline (Cz) and centro-parietal midline (CPz). ERPs are shown during hits, during correct rejections (CRs), and response-locked false alarms (FAs), for young participants of the 1-back Go/NoGo group (panel **A**) and the 2-back Go/NoGo group (panel **B**). The shaded regions around the ERP traces indicate the Standard Error of the Mean (SEM) across participants of the same inhibitory load group.

#### Correct Rejections

Walking-related changes in neural activity during correct rejections were tested for correlations with inhibitory load, independently of d’ score during sitting. The entire set of 64 electrodes and all the epoch timepoints were examined for the existence of such correlations. Walking-*minus*-sitting mean ERP amplitude during correct rejections was calculated at each of the 64 electrodes and each epoch timepoint and it was subsequently subjected to pointwise partial Pearson correlations with inhibitory load (1-back, 2-back), controlling for sitting d’ score. Cluster-based permutation tests were used to identify spatiotemporal clusters of significant correlation effects while accounting for multiple electrode/timepoint tests. During correct rejection trials, walking-*minus*-sitting mean ERP amplitudes were found to negatively correlate with inhibitory load over frontal scalp regions, from 140 ms to 520 ms post-stimulus-onset approximately. These walking-related effects are represented by the blue cluster in the Fig. 5A statistical clusterplot (within-cluster mean Pearson’s r = −0.23, p = 0.0456). Since this cluster spanned multiple stages of inhibitory processing, in order to facilitate its study, six 30-ms time windows were defined such that they were evenly distributed throughout the [140 ms, 520 ms] interval with a distance of 60 ms from each other. The centers of the six 30-ms windows that occurred based on this definition are the following: 180 ms (window 1), 240 ms (window 2), 300 ms (window 3), 360 ms (window 4), 420 ms (window 5), 480 ms (window 6). As inhibitory load increased while walking, walking-related ERP amplitudes were found to become more negative over midline frontocentral regions during windows 1-3, as well as over midline and right-lateralized prefrontal regions during windows 2-6 (Fig. 5A). Fig. 5B shows sitting and walking ERP waveforms during correct rejection trials for each load group at Fpz (frontopolar midline), since this electrode exhibited significant effects for 4 out of the 6 windows (Fig. 5B). The six 30-ms windows are depicted as gray bars on the ERP plots. The topographical maps of Fig. 5C illustrate the scalp distribution of the walking-*minus*-sitting mean ERP amplitudes during correct rejection trials, averaged within each one of the six 30-ms windows separately, for 1-back and 2-back participants.

**Fig. 5.**
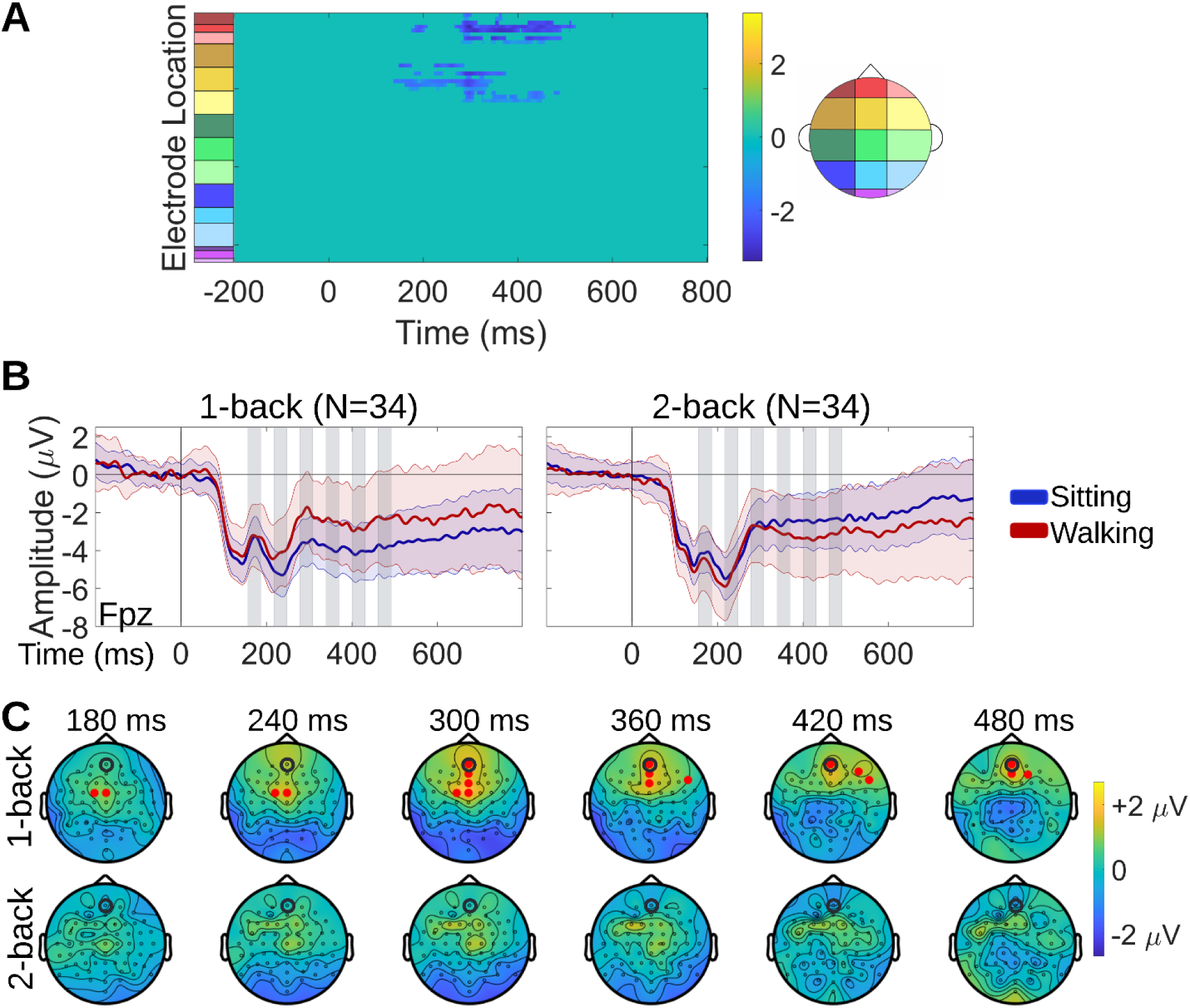
Correlation effects between dual-task-related neural activity change and inhibitory load during correct rejection trials. **A.** Spatiotemporal clusters of walking-*minus*-sitting mean ERP amplitudes correlating with inhibitory load during correct rejections, identified using cluster-based permutation tests. The statistical clusterplot shows the t-values for the electrode-timepoint pairs at which significant correlation was found. Negative t-values (blue) indicate that, as inhibitory load increased while walking, walking ERP amplitudes became more negative relative to the sitting ERP amplitudes. Such significant correlation effects were found over midline frontocentral scalp during ~140-330 ms, and over midline and right-lateralized prefrontal scalp during ~270-520 ms post-stimulus-onset. **B.** Grand mean sitting and walking ERP waveforms for the 1-back and the 2-back Go/NoGo group, at a midline frontopolar electrode (Fpz) that exhibited significant correlation effects. **C.** Topographical maps showing the scalp distribution of the walking-*minus*-sitting mean ERP amplitudes during correct rejections, averaged within each of the load groups separately, for six 30-ms windows (gray bars on the Fig. 5B ERP plots). For each window, the electrodes that exhibited significant correlation effects are indicated by red dots on the topographical map of the load group that was found to drive the correlation. The correlation was driven by the 1-back Go/NoGo group for all six windows. The Fpz electrode to which the ERP waveforms correspond is circled in black on the maps.

Follow-up tests were performed to determine whether the correlation effects detected in each of the windows were driven by a specific load group. Specifically, for each window and within each load group, the mean ERP amplitude, averaged across electrode-timepoint pairs belonging to the respective portion of the significant blue cluster, was compared between sitting and walking using paired t-tests (or Wilcoxon signed rank tests, in case of non-normally distributed data). To correct for the multiplicity of the tests conducted here, specifically (2 load levels)x(6 windows) = 12 tests, the 2-stage step-up False Discovery Rate (FDR) method [84] was applied to adjust the significance level. Significant ERP amplitude differences between sitting and walking were found in all 6 defined windows for 1-back participants (p_window-1_ = 0.0071, p_window-2_ = 0.0007, p_window-3_ < 0.0001, p_window-4_ < 0.0001, p_window-5_ = 0.0009, p_window-6_ = 0.0041; *α_FDR_* = 0.0071), but in none of the windows for 2-back participants (p_window-1_ = 0.3974, p_window-2_ = 0.1347, p_window-3_ = 0.2302, p_window-4_ = 0.9593, p_window-5_ = 0.9251, p_window-6_ = 0.5938; *α_FDR_* = 0.0071). These results indicate that correlation effects were driven by significantly less negative ERP amplitudes when walking under 1-back Go/NoGo load, for all time windows. For each window, the electrodes that exhibited significant correlation effects are denoted by red dots on the topographical map of the load group that was found to drive the correlation (Fig. 5C). The correlation was found to be driven by the 1-back Go/NoGo group for all 6 windows.

These data indicate that attenuation in walking-related ERP amplitude changes over frontal scalp regions, extending from 140 ms to 520 ms post-stimulus-onset approximately, constitute neural signatures of increased inhibitory load in correct rejection trials.

#### Hits

Walking-related changes in neural activity during hits were tested for correlations with inhibitory load, independently of d’ score during sitting, across the entire 64-electrode set and all the epoch timepoints. Walking-*minus*-sitting mean ERP amplitude during hits was calculated at each of the 64 electrodes and each epoch timepoint and it was subsequently subjected to partial Pearson correlations with inhibitory load (1-back, 2-back), controlling for sitting d’ score. Cluster-based permutation tests were used to identify spatiotemporal clusters of significant correlation effects while accounting for multiple electrode/timepoint tests. During hit trials, walking-*minus*-sitting mean ERP amplitudes were found to positively correlate with inhibitory load over predominantly left-lateralized parietal scalp regions, from 50 ms to 470 ms post-stimulus-onset approximately. These walking-related effects are represented by the yellow cluster in the Fig. 6A statistical clusterplot (within-cluster mean Pearson’s r = 0.23, p = 0.0458). Since this cluster spanned multiple stages of information processing, in order to facilitate its study, seven 30-ms time windows were defined such that they were evenly distributed throughout the [50 ms, 470 ms] interval with a distance of 60 ms from each other. The centers of the seven 30-ms windows that occurred based on this definition are the following: 80 ms (window 1), 140 ms (window 2), 200 ms (window 3), 260 ms (window 4), 320 ms (window 5), 380 ms (window 6), 440 (window 7). As inhibitory load increased while walking, walking-related ERP amplitudes were found to become more positive over left parietal regions during windows 1-5, and over central regions during windows 6-7 (Fig. 6A). Fig. 6B shows sitting and walking ERP waveforms during hit trials for each load group at electrode P3 (left parietal), since this electrode exhibited significant effects for 5 out of the 7 windows (Fig. 6B). The seven 30-ms windows are depicted as gray bars on the ERP plots. The topographical maps of Fig. 6C illustrate the scalp distribution of the walking-*minus*-sitting mean ERP amplitudes during hit trials, averaged within each one of the seven 30-ms windows separately, for 1-back and 2-back participants.

**Fig. 6.**
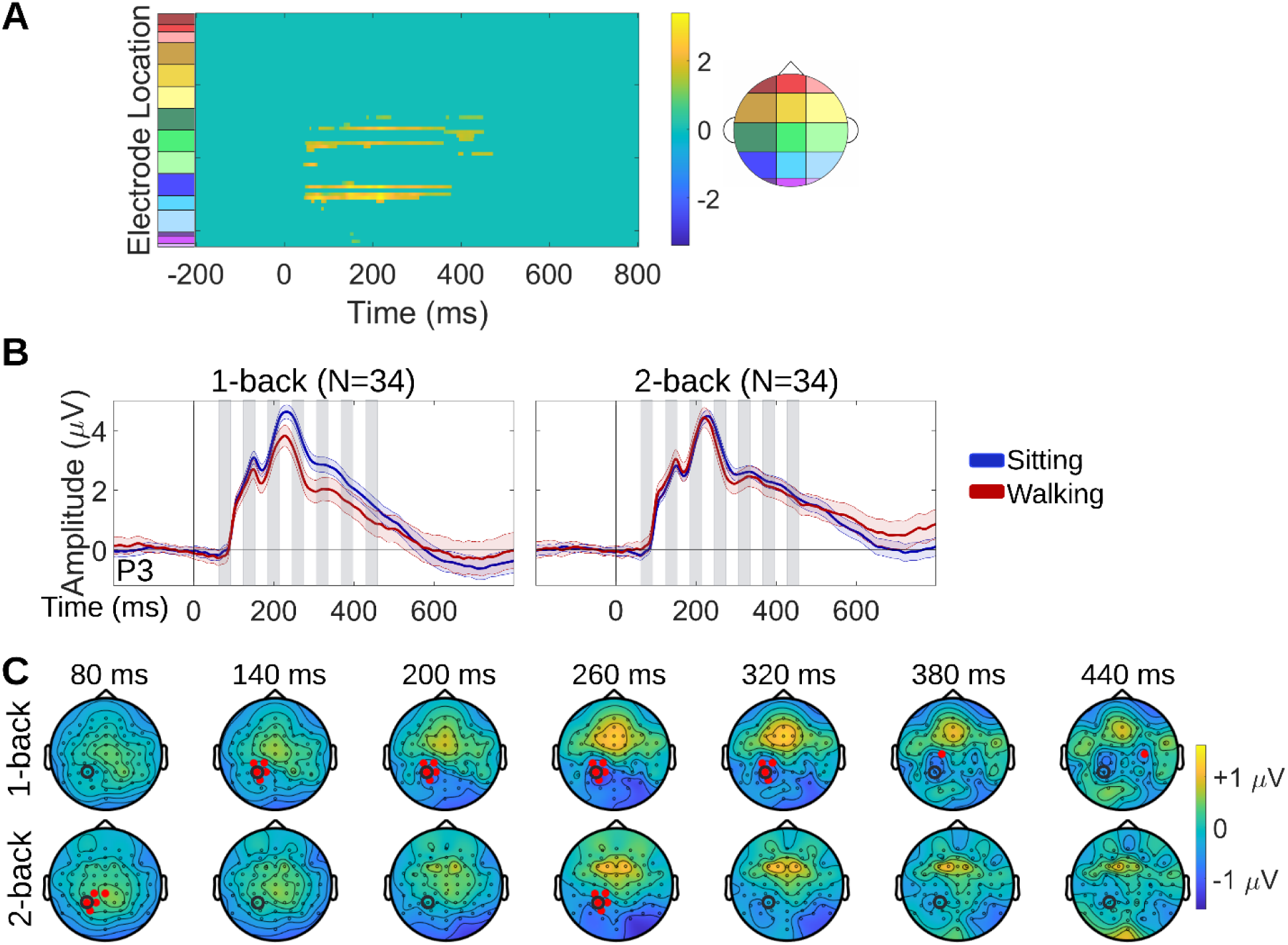
Correlation effects between dual-task-related neural activity change and inhibitory load during hit trials. **A.** Spatiotemporal clusters of walking-*minus*-sitting mean ERP amplitudes correlating with inhibitory load during hits, identified using cluster-based permutation tests. The statistical clusterplot shows the t-values for the electrode-timepoint pairs at which significant correlation was found. Positive t-values (yellow) indicate that, as inhibitory load increased while walking, walking ERP amplitudes became more positive relative to the sitting ERP amplitudes. Such significant correlation effects were found over left-parietal scalp during ~50-350 ms, and over central scalp during ~350-470 ms post-stimulus-onset. **B.** Grand mean sitting and walking ERP waveforms for the 1-back and the 2-back Go/NoGo group, at a left parietal electrode (P3) that exhibited significant correlation effects. **C.** Topographical maps showing the scalp distribution of the walking-*minus*-sitting mean ERP amplitudes during hits, averaged within each of the load groups separately, for seven 30-ms windows (gray bars on the Fig. 6B ERP plots). For each window, the electrodes that exhibited significant correlation effects are indicated by red dots on the topographical map of the load group that was found to drive the correlation. The correlation was driven by the 1-back Go/NoGo group for windows 2-7, and by the 2-back Go/NoGo group only for windows 1 and 4. The P3 electrode to which the ERP waveforms correspond is circled in black on the maps.

Follow-up tests were performed to determine whether the correlation effects detected in each of the windows were driven by a specific load group. Specifically, for each window and within each load group, the mean ERP amplitude, averaged across electrode-timepoint pairs belonging to the respective portion of the significant yellow cluster, was compared between sitting and walking using paired t-tests (or Wilcoxon signed rank tests, in case of non-normally distributed data). To correct for the multiplicity of the tests conducted here, specifically (2 load levels)x(7 windows) = 14 tests, the significance level was adjusted using the 2-stage step-up FDR method. In the 1-back Go/NoGo group, significant ERP amplitude differences between sitting and walking were found in windows 2-7 (p_window-1_ = 0.6552, p_window-2_ = 0.0032, p_window-3_ = 0.0039, p_window-4_ < 0.0001, p_window-5_ = 0.0002, p_window-6_ = 0.0148, p_window-7_ = 0.0029; *α_FDR_* = 0.0148). In the 2-back Go/NoGo group, such differences were found only in windows 1 and 4 (p_window-1_ < 0.0001, p_window-2_ = 0.3604, p_window-3_ = 0.1236, p_window-4_ = 0.0009, p_window-5_ = 0.1990, p_window-6_ = 0.3341, p_window-7_ = 0.0990; *α_FDR_* = 0.0148). These results indicate that correlation effects were driven by significantly more positive ERP amplitudes when walking under 2-back Go/NoGo load for window 1, and by significantly less positive ERP amplitudes when walking under 1-back Go/NoGo load for all the rest of the windows (2-7). For each window, the electrodes that exhibited significant correlation effects are denoted by red dots on the topographical map of the load group that was found to drive the correlation (Fig. 6C). As this figure shows, along with Fig. 6B, the correlation was found to be driven by the 1-back Go/NoGo group for windows 2-7, and by the 2-back Go/NoGo group only for windows 1 and 4.

Based on these data, increased walking-related amplitudes over left parietal regions during 50-110 ms post-stimulus-onset approximately, and attenuated walking-related amplitude changes over left parietal regions during 110-350 ms and over central regions during 350-470 ms approximately, constitute neural signatures of increased inhibitory load in hit trials.

### Gait

Fig. 7B demonstrates the stride-to-stride variability distributions with and without a concurrent cognitive task (WT and WO, respectively), with the cognitive task being either 1-back Go/NoGo or 2-back Go/NoGo. Fig. 7A illustrates the 3D representations of trajectories of a series of strides for one young participant while performing the 1-back Go/NoGo task, and one young participant while performing the 2-back Go/NoGo task (panel A).

**Fig. 7.**
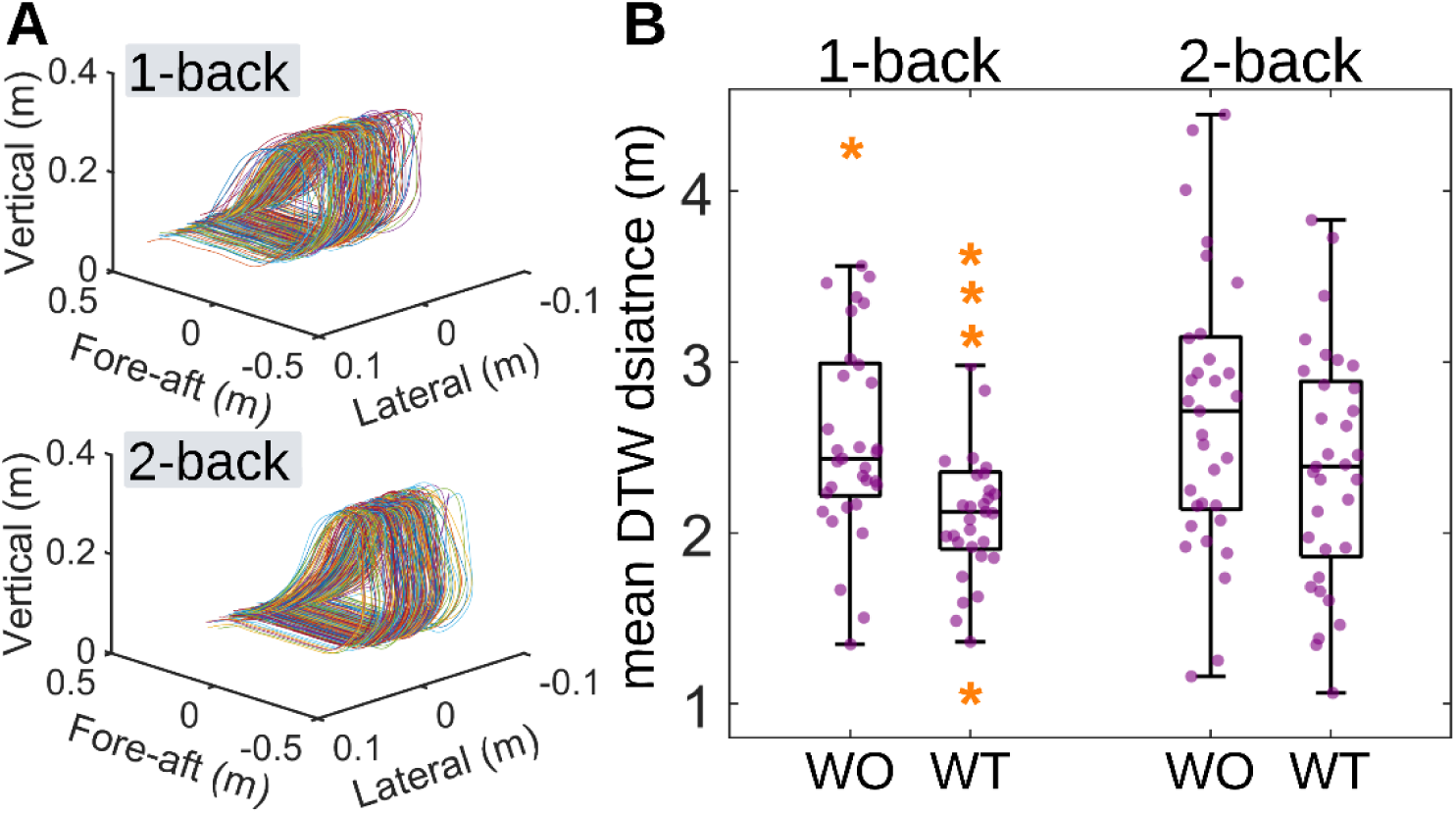
**A.** 3D representations of trajectories of a series of strides for one young participant while engaging in the 1-back Go/NoGo task, and one young participant while engaging in the 2-back Go/NoGo task. Lateral is the dimension of movement right-and-left relative to the motion of the treadmill belt. Vertical is the dimension movement up-and-down relative to the motion of the treadmill belt. Fore-aft is the dimension of movement parallel to the motion of the treadmill belt. Using Dynamic Time Warping (DTW), the variability from one stride to the next was quantified as DTW distance (see Methods) and the mean DTW distance of all stride-to-stride comparisons was extracted per participant. **B.** Mean DTW distance distributions during walking with task (WT) and walking only (WO), for the 1-back Go/NoGo group and the 2-back Go/NoGo group. Purple dots represent individual participants. Orange asterisks represent outliers.

Potential associations were tested between WT-minus-WO mean DTW distance and inhibitory load, controlling for sitting d’ score, using a partial Pearson rank correlation. The correlation was found to be non-significant (Pearson’s r = 0.02, p = 0.8929), thereby indicating that increasing inhibitory load from 1-back to 2-back Go/NoGo while walking has no detectable effect on stride-to-stride variability.

## DISCUSSION

As inhibitory load increased from 1-back to 2-back Go/NoGo, response accuracy during sitting declined. This result could be expected to some extent based on previous studies which observed deterioration of response accuracy when working memory load was increased from 1-back to 2-back [21, 85]. However, the inhibitory component was not present in the task designs of those studies, so the present finding of response accuracy decline with increasing inhibitory load is novel. Regarding the addition of walking to cognitive task performance, it was found to improve response accuracy in most young adults at both the lower (1-back) and the higher (2-back) inhibitory load level, specifically in 20 out of 34 participants in both cohorts. Also, no detectable difference in walking-related response accuracy changes was found between the two load levels. Although this walking-related improvement contradicts the CMI hypothesis, it has been reported before, when pairing walking with the 1-back Go/NoGo task in young adults [10]; as such, it was anticipated for the 1-back Go/NoGo load level. This improvement has been attributed to increased arousal putatively induced by moderate forms of exercise, such as walking [86–89] (see Patelaki and colleagues [10, 55] for a more detailed explanation). However, contrary to what was hypothesized, young adults managed to maintain response accuracy improvement during walking even under increased, 2-back Go/NoGo load. This maintained improvement in terms of response accuracy was not accompanied by costs in any other behavioral aspect, such as response speed or gait consistency, but it came with walking-related ERP amplitude changes during correct rejections and hits.

During correct rejection trials, as inhibitory load increased while walking, walking-related ERP amplitudes increased over midline frontocentral regions during 140-330 ms approximately, as well as over midline and right-lateralized prefrontal regions during 270-520 ms approximately (Fig. 5A). These effects were driven by attenuation in walking-related ERP amplitude changes under 2-back Go/NoGo load (Figs. 5B and 5C).

The frontocentral portion of walking-related effects during 140-330 ms, post-stimulus-onset, of correct rejection trials is associated with the N1 and the N2 stages of inhibitory processing. The N1 is the negative voltage deflection peaking frontocentrally ~100-200 ms post-stimulus-onset [90–94], and it is thought to reflect sensory gain control processes, involving suppressing irrelevant stimulus information and allocating top-down attentional resources towards amplifying processing of relevant stimulus information [24, 94–97]. The N2 is the negative voltage deflection peaking frontocentrally ~200-350 ms post-stimulus-onset [32–34] and it reflects conflict monitoring processes [35–37, 79]. Reductions in N1 and N2 amplitudes during walking have been previously found in IMPs [10, 55], and have been interpreted as markers of flexible reallocation of neural resources when dual-task walking. Conversely, young NCs and DECs, who presumably lack such flexibility, showed absence of walking-related reductions in terms of these ERP components. Based on these previous findings, the absence of walking-related N1 and N2 reductions as inhibitory load increases while walking, which was exhibited here by the 2-back Go/NoGo group, likely reflects more effortful reallocation of neural resources during processing related to sensory gain control and conflict monitoring.

The more anterior (midline and right-lateralized prefrontal) portion of walking-related effects during 270-520 ms, post-stimulus-onset, of correct rejection trials is related to the N2 and the P3 stages of inhibitory processing. During the P3 stage, which takes place ~350-600 ms post-stimulus-onset [6, 38], neural processes underpinning both the motor and the cognitive component of response inhibition are thought to be executed [39, 40]. Part of the processing related to the ‘higher-level’ cognitive component of inhibition is related to appropriately maintaining and updating the task representations so that behavioral performance in ensuing trials is successful. Given that the 2-back Go/NoGo task requires keeping task representations in the memory buffer for a sequence of preceding trials ‒ and not just for the immediately previous trial, as is the case with the 1-back Go/NoGo task ‒ additional recruitment of neural resources related to working memory is expected under 2-back Go/NoGo load. Anterior prefrontal regions, such as the right-lateralized frontopolar cortex, have been found to play a special role in maintaining, manipulating and updating abstract, internal representations subserving a secondary or concurrent task goal in the context of a complex task implicating working memory. Specifically, when integrating working memory with a second executive function in the same task, the occurring higher-level, integration-related representations are presumably stored and manipulated in the frontopolar cortex [98–101], with existing evidence particularly emphasizing the role of its right-hemisphere portion [98]. In the 2-back Go/NoGo task implementation employed here, which combines working memory with response inhibition, these dedicated working memory resources of the frontopolar cortex presumably serve to maintain, manipulate and update inhibition-related internal representations, in order to achieve successful performance in 2-back response inhibition. In the 1-back Go/NoGo group, walking-related changes in ERP amplitudes during the N2 and the P3 stages over anterior prefrontal regions likely reflect their flexible recruitment when dual-task walking. On the other hand, in the 2-back Go/NoGo group, attenuation of these ERP changes indicates that 2-back Go/NoGo load likely brings these working memory storage and processing resources so close to their tipping point that the addition of walking has little to no impact on their reallocation.

During hit trials, as inhibitory load increased while walking, walking-related ERP amplitudes increased over left parietal regions during 50-350 ms approximately, and over central regions during intervals 350-470 ms approximately (Fig. 6A). These effects were driven predominantly by attenuation in walking-related ERP amplitude changes under 2-back Go/NoGo load. (Figs. 6B and 6C)

The left parietal portion of walking-related effects during ~50-350 ms, post-stimulus-onset, of hit trials spanned multiple stages of information processing, from sensory and perceptual to response selection [90, 102]. The observation that these effects were not associated to a specific processing stage indicates that they likely reflect walking-related changes in ‘slow’ neural processes housed in the left parietal cortex. The left parietal cortex has been shown to play a key role in motor attentional control [103, 104], such as in orienting attention to and selecting the appropriate motor plan which, in the case of hits, is generating a response by pressing the button. Attenuations in walking-related ERP amplitude changes over left-dominant parietal regions have been previously found in older adults, during the N2 stage of correct rejections [55]. These aging-related ERP effects have been interpreted as markers of reduced flexibility in reallocating motor-attentional resources under conditions of increased task demands. By extension, the attenuated walking-related ERP amplitude changes that young adults of the 2-back Go/NoGo group exhibited left-parietally, and during the N1 and N2 stages of hits (~110-350 ms post-stimulus-onset), might reflect less flexible, or less automatic, reallocation of motor-attentional resources as inhibitory load increases while walking. Even though these load-related parietal signatures were manifested under a different behavioral condition (hits) compared to the corresponding aging-related parietal signatures (correct rejections), they are possibly linked to the same underlying motor-attentional process, since preparedness to update the motor plan according to the rapidly changing task needs is necessary across trials and regardless of behavioral condition. Regarding the early (~50-110 ms) walking-related increase in left-parietal ERP amplitudes which was observed under 2-back Go/NoGo load, but was absent under 1-back Go/NoGo load, it should be interpreted with caution because of the small size of the effect. Specifically, this load-related effect had a short duration, and the walking-*minus*-sitting ERP amplitude associated with it was very small (Fig. 6B). If a speculation about it must be provided, this early walking-related ERP amplitude increase under 2-back Go/NoGo load might reflect an effort to urgently recruit the left-parietal resources before they reach their tipping point later in the processing stream and, in that way, to avoid performance decline during walking.

The central portion of walking-related effects during ~350-470 ms, post-stimulus-onset, of hit trials corresponds to the P3 component. For hits, P3 reflects neural processes related to the execution of the motor response [39, 41, 105, 106]. These processes have been traced to central—such as motor and mid-cingulate—cortical regions, aligning with the scalp topography of the walking-related neural activity effects during the latencies of interest (Figs. 6A and 6C) [41, 44, 105–108]. When exposed to 2-back Go/NoGo load, young adults exhibited attenuations in walking-related ERP amplitude changes over these central topographies. Similar to the parietal effects above, attenuations in walking-related ERP amplitude changes over central regions have been previously found in older adults, during the P3 stage of correct rejections [55]. These aging-related ERP effects are thought to reflect reduced flexibility in reallocating motor-executional resources under conditions of increased task demands. Despite the difference in the behavioral condition during which attenuation effects were detected (hits for 2-back versus correct rejections for aging), central cortical regions are putatively implicated in the execution of the appropriate motor plan, both when this is a ‘Go’ and when this is a ‘NoGo’ [39, 41, 42, 44, 105, 106]. Extending this interpretation, the attenuation in walking-related ERP amplitude changes observed in the 2-back Go/NoGo group centrally, and during the P3 stage of hits, were interpreted as load-related reduction of automaticity in recalibrating neural processes related to executing the ‘Go’ button press when dual-task walking.

As discussed in the paragraphs above, increasing inhibitory load from 1-back to 2-back Go/NoGo during walking comes with attenuations in walking-related ERP amplitude changes, a pattern that, interestingly, has been previously encountered in aging, when walking under 1-back Go/NoGo load. Despite this similarity, there are also differences between the neurophysiological effects of 2-back Go/NoGo load and aging during dual-task walking, specifically in terms of their scalp topographical distribution. During correct rejections, aging-related attenuations were detected over left frontal, left parieto-occipital and centro-parietal regions [55], while load-related attenuations were found herein over frontocentral and right prefrontal regions. Furthermore, in terms of behavioral performance, increased inhibitory load was associated with no detectable d’ change during walking, but aging has been previously found to come with walking-related d’ decline [30, 55]. Comparative assessment of these findings suggests that, even though both increased inhibitory load and aging indeed tax the neural resources during dual-task walking, the brain areas and processes sustaining the tax are different in each case. Also, based on the behavioral findings, aging likely taxes the neural resources beyond their tipping point during dual-task walking, while 2-back Go/NoGo load seems to ‘push’ the neural resources closer but not past their tipping point in younger adults.

One limitation of this study is that the 1-back Go/NoGo group and the 2-back Go/NoGo group consisted mostly of different young adults, with an overlap of only 7 people who participated in both groups. Recruitment for the 2-back cohort started more than a year later than recruitment for the 1-back cohort. This gap resulted in high attrition rates, since our 1-back cohort consisted primarily of university students who had graduated and moved away from campus by the time recruitment for the 2-back cohort started, and as such they were no longer available or interested to participate. To minimize any potential unwanted inter-group variability, future studies are recommended to recruit the same individuals to participate in both dual-tasks in two separate visits, ideally with a gap of no more than a few weeks between the two visits. Also, it should be noted that, extracting the approximate onset and offset latencies of the detected spatiotemporal clusters was only utilized to estimate the processing stages that the interrogated correlation effects spanned (e.g., N1, N2, P3, ERN). Interpretation of the precise onset and offset latencies of the clusters was avoided since that would be an incorrect use of the cluster-based permutation testing approach [109].

## CONCLUSIONS

In summary, under conditions of concurrent walking and Go/NoGo response inhibition task performance, young adults maintained improvement in response accuracy even when the inhibitory load imposed by the task was increased. Increased inhibitory load while walking was not accompanied by any detectable costs in gait consistency or response speed, but it was found to correlate with attenuations in walking-related ERP amplitude changes during correct rejections and hits. In particular, during correct rejections, such attenuations were detected over frontal regions, during latencies related to sensory gain control, conflict monitoring and working memory storage and processing. During hits, attenuations were found over left-parietal regions, during latencies related to orienting attention to and selecting the ‘Go’ motor plan, and also over central regions, during latencies linked to executing the ‘Go’ motor response. The pattern of attenuation in walking-related ERP amplitude changes in the 2-back Go/NoGo group was interpreted as a manifestation of increased effort to recalibrate the above neural processes when pairing walking with increased inhibitory load. This widespread and more effortful reallocation of neural resources is likely an effective mechanism working to ‘shield’ against performance costs when inhibitory load increases during walking. Activating and utilizing such a mechanism suggests that increasing inhibitory load during walking likely brings the neural resources of young adults closer to their tipping point, but not past it. Determining the tipping point of the neural resources of young adults can help assess the neurocognitive bandwidth that young, flexible brains possess. Such information could hold potential for better understanding how this bandwidth gets curtailed in aging or neurological disorders.

## AUTHOR CONTRIBUTIONS

EP: Conceptualization, Data Curation, Formal Analysis, Investigation, Methodology, Software, Visualization, Writing – Original Draft Preparation; JJF: Conceptualization, Funding Acquisition, Methodology, Project Administration, Supervision, Writing – Original Draft Preparation; ALM: Investigation, Writing - Review & Editing; EGF: Conceptualization, Funding Acquisition, Methodology, Project Administration, Supervision, Writing – Original Draft Preparation

## ETHICS STATEMENT

This study was conducted in accordance with the tenets of the Declaration of Helsinki and with approval from the Institutional Review Board of the University of Rochester (STUDY00001952). All participants provided written informed consent.

## DATA/CODE AVAILABILITY STATEMENT

Data from this study will be made available through a public repository (e.g. Dryad) upon publication of this paper, and the authors will work with the editorial office during production to incorporate appropriate links. Custom code from this study will be made available on GitHub (https://github.com/CNL-R) upon publication of this paper.

## Abbreviations List

3D: three-dimensional
CMI: cognitive motor interference
CR: correct rejection
DTW: dynamic time warping
EEG: electroencephalography
ERN: error-related negativity
ERP: event-related potential
MoBI: mobile brain-body imaging
IMPs: participants who improved when dual-task walking
NCs: participants whose performance did not change between sitting and walking
DECs: participants who declined when dual-task walking
RT: response time
WT: walking with task
WO: walking only

## ACKNOWLEDGEMENTS

We would like to thank each of the participants that enrolled in the study.

